# Division of vocal signaling in cooperating marmoset dyads

**DOI:** 10.64898/2026.10.15.758668

**Authors:** Gargi Nandy, Olivia C. Meisner, Weikang Shi, Anirvan S. Nandy, Monika P. Jadi, Steve W. C. Chang

**Author notes:** co-senior authors (Anirvan S. Nandy, Monika P. Jadi).

## Abstract

Communication enables partners to coordinate actions toward shared goals, but how vocal communication is organized during cooperation remains poorly understood. We recorded vocalizations from three common marmoset (*Callithrix jacchus*) dyads as they learned a mutual cooperation task requiring coordinated lever pulls for reward. Vocal behavior was divided by social hierarchy: dominants produced t-peeps, whereas subordinates produced chirps. Interestingly, dominant t-peeps followed failed cooperative pulls, peaked when cooperative bouts broke down, and declined after learning. By contrast, subordinate chirps followed successful pulls, accumulated during sustained cooperation, and persisted after learning. Each call was linked to the caller’s task role: dominant animals t-peeped more in sessions they led, and subordinates chirped mainly after cooperations they initiated, associations that sharpened with learning. Thus, social rank determined which vocal signal a partner used, whereas task role and the state of cooperation determined when to communicate. These complementary vocal signals of failure and success reveal how hierarchy, task role and task phase organize communication during joint action in cooperating marmosets.

## INTRODUCTION

Communication allows social partners to coordinate their behavior toward shared goals. Common marmosets (*Callithrix jacchus*) have emerged as a powerful nonhuman primate model for studying the biological foundations of such interactive communication^1^. As cooperative breeders, marmosets are highly prosocial, possess a rich vocal repertoire, and engage in reciprocal vocal exchanges characterized by turn-taking and sensitivity to social feedback^2–5^. Their vocal behavior is also flexible: social experience shapes vocal development and ongoing interactions influence when and how animals call^3,4,6,7^. Together with mutual gaze and a strong propensity for cooperation, these features have led marmosets to be proposed as a model for the social interactional foundations of human communication^5,6,8,9^. Yet how vocal communication is organized when two individuals must coordinate their actions to achieve a shared goal remains poorly understood.

An important setting to address this question is cooperation in these species, where successful behavior depends explicitly on coordinating actions with another individual. We have recently shown that marmoset dyads learn to work with one another in an automated lever-pulling task in which both partners must pull their own levers within a short temporal window to obtain mutual rewards10. Importantly, marmosets did not solve this task using a single stereotyped strategy. Instead, dyads developed flexible, partner-dependent strategies involving social monitoring and coordination, and these strategies were shaped by dominance, kinship, and sex. In particular, dominant animals engaged in greater social gaze during periods of unsuccessful coordination and learned to adjust their pulling behavior relative to their partners, suggesting that the two members of a dyad can assume differentiated roles during the acquisition of cooperation. Given that vocal communication is a defining feature of this species’ social interactions, we hypothesized that, like social gaze monitoring, it is also used when they learn to cooperate by contributing to their partner-dependent behavioral strategies.

Here, we asked whether vocal communication is strategically used and, if so, how vocal behaviors are organized between social partners as they learn to cooperate. We recorded vocalizations from three familiar marmoset dyads during acquisition and subsequent performance of the mutual cooperation task. We examined how call production depended on social rank, learning stage, the outcome of individual cooperative attempts, the partner who initiated each cooperation, and the moment-to-moment state of cooperation within a session. We found that of four common call types (phees, trills, t-peeps and chirps), only t-peeps and chirps were produced at appreciable rates during the mutual cooperation task, and the uses of these calls were strongly dependent on the rank of the vocalizing animal. Dominant animals produced t-peeps after failed cooperative attempts and at the breakdown of cooperative bouts, and these calls declined once the task was learned. By contrast, subordinate animals produced chirps after successful cooperative pulls and throughout sustained cooperation, and these calls persisted after learning. Each call was further linked to preferential usage during the caller’s own leadership of the cooperation. Finally, the partners vocalized preferentially within and outside cooperative bouts in a session. These findings reveal that vocal communication during mutual cooperation can be distributed asymmetrically across social partners, with different members of a dyad providing complementary signals about the outcome of their joint behavior.

## RESULTS

We recorded vocalizations from three familiar dyads as they learned to pull their levers to receive mutual rewards in an automated cooperative pulling task^10,11^. Each marmoset was placed in its own transparent chamber with a lever, and both earned a liquid reward only when the two animals pulled within 1 s of each other (Figure 1A). Vocal behaviors from each dyad were recorded during the “training’ stage and the ‘trained’ stage. These two stages were distinguished by either success rate or repeated performance: the dyad had to either reach a 50% success rate or perform during four sessions of the given condition during training (see Methods) before moving to the trained stage.

**Figure 1.**
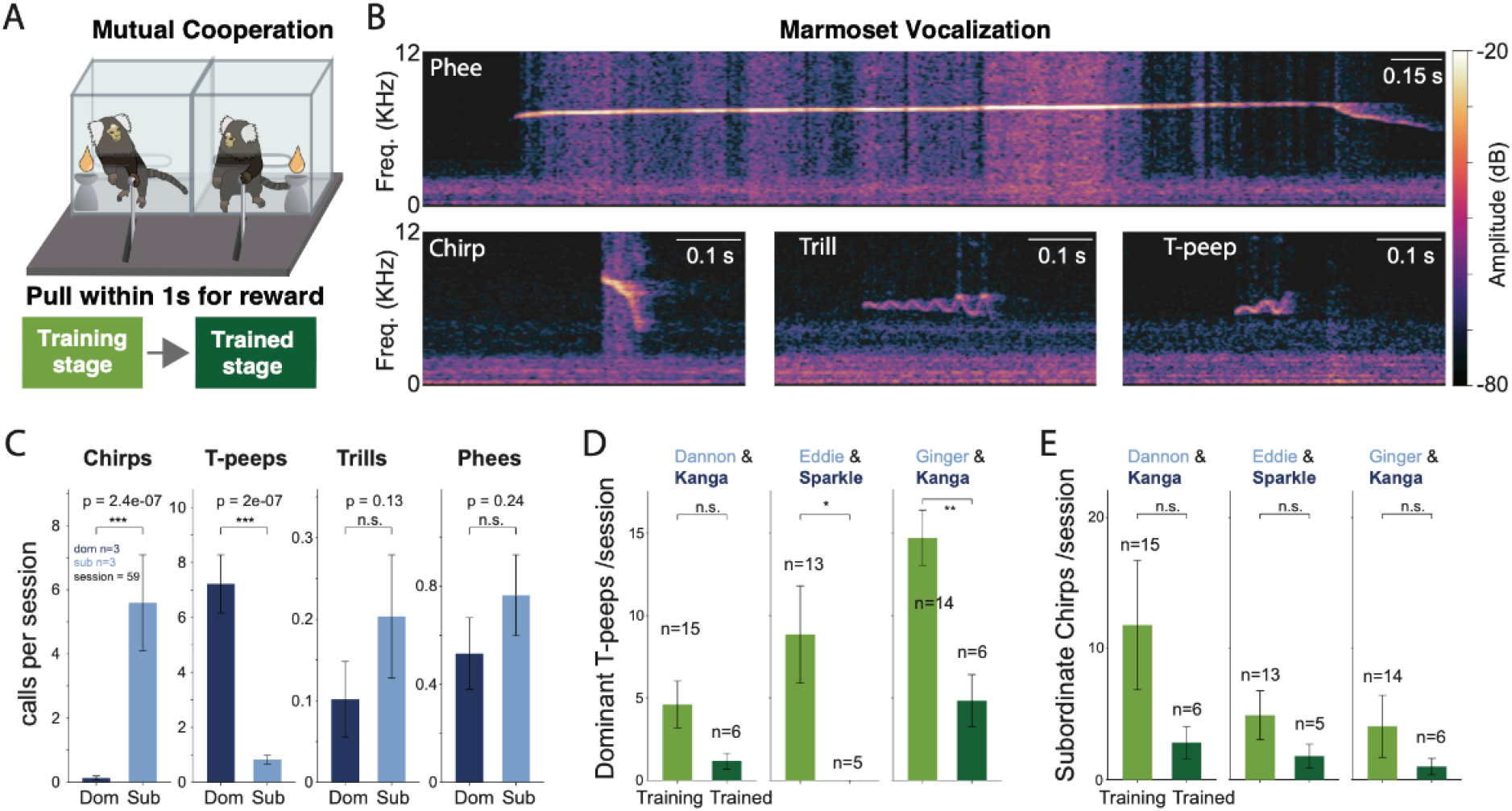
Vocalization call types during cooperation as a function of social hierarchy. (A) The mutual cooperation task. Two marmosets sit in adjacent transparent chambers, each with its own lever. Both are rewarded only when both pull their levers within 1 s of each other. Each dyad was recorded during the Training stage (light green) and again once trained (Trained stage, dark green). (B) Example spectrograms of the four call types analyzed: phee, chirp, trill and T-peep. The color scale shows amplitude in dB. See Supplementary Information for more examples.(C) Calls per session by the dominant (Dom, dark blue) and subordinate (Sub, light blue) monkey for each call type. Data come from both Training and Trained sessions with all three dyads pooled (n = 59 sessions; 3 dominant and 3 subordinate monkeys per dyad role). (D) The dominant monkey’s t-peeps per session in the Training and Trained stages, shown for each dyad. The dominant is signified by dark blue. (E) Same as (D) for the subordinate monkey’s chirps. In C–E, bars are mean ± SEM across sessions and n is the number of sessions. C uses paired Wilcoxon signed-rank tests across sessions; D and E use Mann–Whitney U tests. ^*^ p < 0.05, ^**^ p < 0.01, ^***^ p < 0.001; n.s., not significant.

We analyzed four call types (Figure 1B): phee, trill, t-peep and chirp^12–17^. Phees are long (0.5–2 s), tonal contact calls usually produced when animals are separated^16^. Trills are low-intensity calls with sinusoidal frequency modulation, used between animals that are in proximity^16^. T-peeps (trill-like peeps) share the trill’s structure but are much shorter (30–200 ms)^16^. Chirps are brief (∼0.05s), sharply descending (10Khz to 6Kz) frequency sweeps that are also classified as food calls in a functional context^13^, and are referred to as sd-peeps when described in terms of their acoustic properties^16^.

### Dominant and subordinate marmosets produce different vocal calls during cooperation training

While the cooperating dyads made vocal calls sparsely, the types of vocal calls produced were strongly dependent on the social rank of the vocalizing animal (Figure 1C). We characterized dominance by observing specific behaviors in the home cage based on established dominance criteria^18^: high-ranking marmosets actively took over food and occupied preferred resting spots using dominant calls and postures, while lower-ranking marmosets submitted by moving away and showing deference^10^. The subordinate animals produced significantly more chirps calls than dominant animals during mutual cooperation (p = 7.8 × 10^−6^, paired Wilcoxon signed-rank test). By contrast, dominant animals produced significantly more t-peeps than subordinates (∼9.2 vs. ∼0.7 calls per session; p = 1.7 × 10^−6^). Trills (∼0.14 vs. ∼0.26; p = 0.2) and phees (∼0.73 vs. ∼0.89; p = 0.5) were rare and did not differ by rank. We therefore focused our further analyses to t-peeps and chirps.

Interestingly, the t-peep rate per session by the dominant animals declined once dyads were trained (Figure 1D). The drop was significant in two dyads (E+S: ∼8.8 to ∼0 per session; n = 13 vs. 5 sessions; p < 0.05, G+K: ∼14.5 to ∼4.8; n = 14 vs. 6; p < 0.01, Mann–Whitney U test) and trending in the same direction for the third dyad (D+K: ∼4.5 to ∼1.2; n = 15 vs. 6; p = 0.597).

While subordinate chirp rate also fell in all dyads from training to trained (D+K ∼12 to ∼3; E+S ∼5 to ∼2; G+K ∼4 to ∼1; all p > 0.072), none of these changes were significant (Figure 1E). Thus, t-peeps, predominantly made by the dominant animals, were tied mainly to the learning period, whereas chirps, almost exclusively made by the subordinate animals, persisted, albeit at lower rates, even after the task was learned.

### Dominant t-peeps follow failed cooperative pulls

We next asked when, relative to the task, dominants produced t-peeps. In the example behavioral time series shown in Figure 2A, the dominant pulled without its partner (a failed cooperative pull), produced a t-peep call, and the dyad then completed a successful cooperative pull on their next attempt. This example series of behavioral events raises an intriguing possibility that the dominant marmoset communicated the failed cooperative attempt to the partner using a t-peep call. When aligned to dominant animals’ pulls, the probability of a dominant t-peep call consistently increased after the dominant’s own failed pulls, but not after their successful ones (Figure 2B, top). In all three dyads, t-peep probability after a failed pull peaked 1–2 s after the pull (Figure 2B, top). By contrast, after successful pulls, the t-peep probability remained near zero after the pull. This pattern could be clearly seen when we only examined the pulls with at least one dominant t-peep within ±5 s of the pulling event (Figure 2C). Within this subset, dominant mostly made t-peeps before a successful pull (up to ∼0.3–0.4 per 1-s bin), but not after the pull succeeded (Figure 2C). After failed pulls, however, dominant animals continued to produce t-peeps (Figure 2C). Dominant marmosets thus emitted t-peeps mainly after failed cooperation, and occasionally up to a successful one.

**Figure 2.**
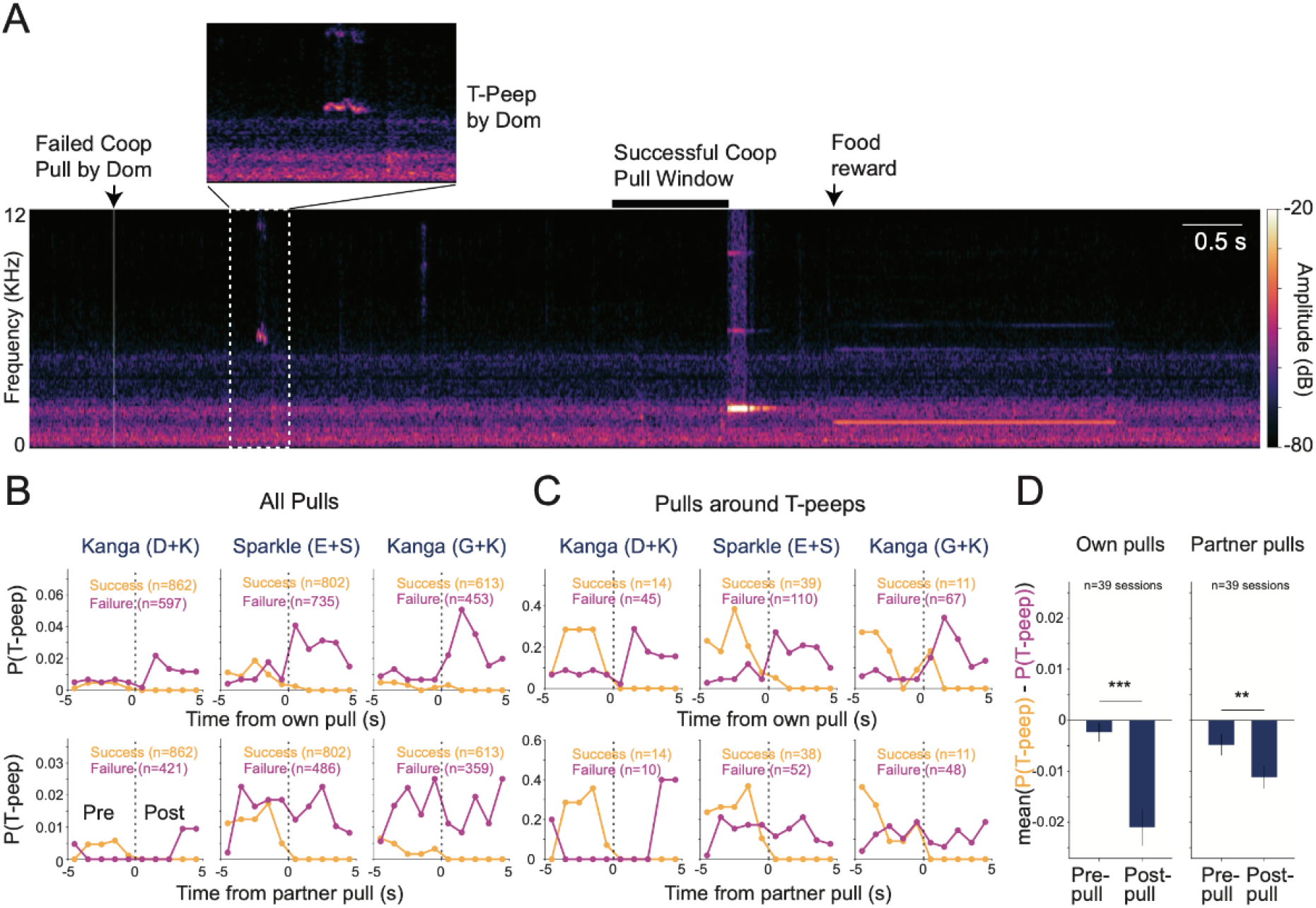
Dominant marmoset’s t-peeps around cooperative pulls. (A) Example spectrogram of a dominant monkey’s audio. It shows a failed cooperative pull by the dominant (arrow), then a T-peep by the dominant (dashed box, enlarged in the inset), then a successful cooperative pull (black bar marks the pull window). The color scale shows amplitude in dB. (B) Probability that the dominant monkey gives at least one T-peep in each 1 s bin, from 5 s before to 5 s after a pull. Pulls are split into successful (orange) and failed (magenta) cooperative pulls; a pull counts as successful when the partner also pulled within 1.0 s. The top row is aligned to the dominant’s own pulls and the bottom row to the partner’s pulls. All pulls from labeled sessions are included; n is the number of pulls. (C) Same as (B), but counting only pulls with at least one of the dominant’s T-peeps within ±5 s. (D) For each session, the mean of P(T-peep | success) − P(T-peep | failure) over the five pre-pull bins and the five post-pull bins, for own pulls (left) and partner pulls (right). Bars show mean ± SEM across sessions (n = 39 sessions, all three dyads pooled). Pre vs. post was compared with a paired Wilcoxon signed-rank test (own pulls p = 5.7 × 105; partner pulls p = 0.0016). ^**^ p < 0.01, ^***^ p < 0.001.

To quantify these cooperation-dependent t-peep patterns by the dominant marmosets across sessions, we compared the average difference in t-peep probability conditioned on success or failure outcome, for both the pre- and post-pull periods (Figure 2D). This difference was near zero prior to the pulls but was significantly negative after the pull, indicating a greater proportion of t-peeps following failed pulls. This pattern was observed for both the dominant’s own pulls (pre ∼−0.002, post ∼−0.021; p = 5.7 × 10^−5^, paired Wilcoxon signed-rank test) and the partner’s pulls (pre ∼−0.005, post ∼−0.011; p = 0.0016). Dominant t-peeps therefore tracked failed cooperation attempts rather than pulling actions in general.

When aligned to the partner’s pulls, t-peeps made by the dominant animals were more common around the partner’s failed pulls than successful pulls, and this probability again decreased to near zero after successful pulls by the partner (Figure 2B, bottom). More importantly, the dominant t-peeps around partner’s failed pull were not as clearly restricted to the post-pull period (Figure 2B, bottom), also reflecting in a smaller change in the pre- and post-pull periods for the difference in conditional probability of t-peeps (Figure 2D, right).

### Subordinate chirps follow successful cooperative pulls

Compared to the dominant marmosets, chirps produced by subordinate marmosets had the opposite pattern. These calls came after successful cooperative pulls. In the example series of behavioral events in Figure 3A, each of the two successful pulls was followed by one or more subordinate chirps that occurred within about 1s.

**Figure 3.**
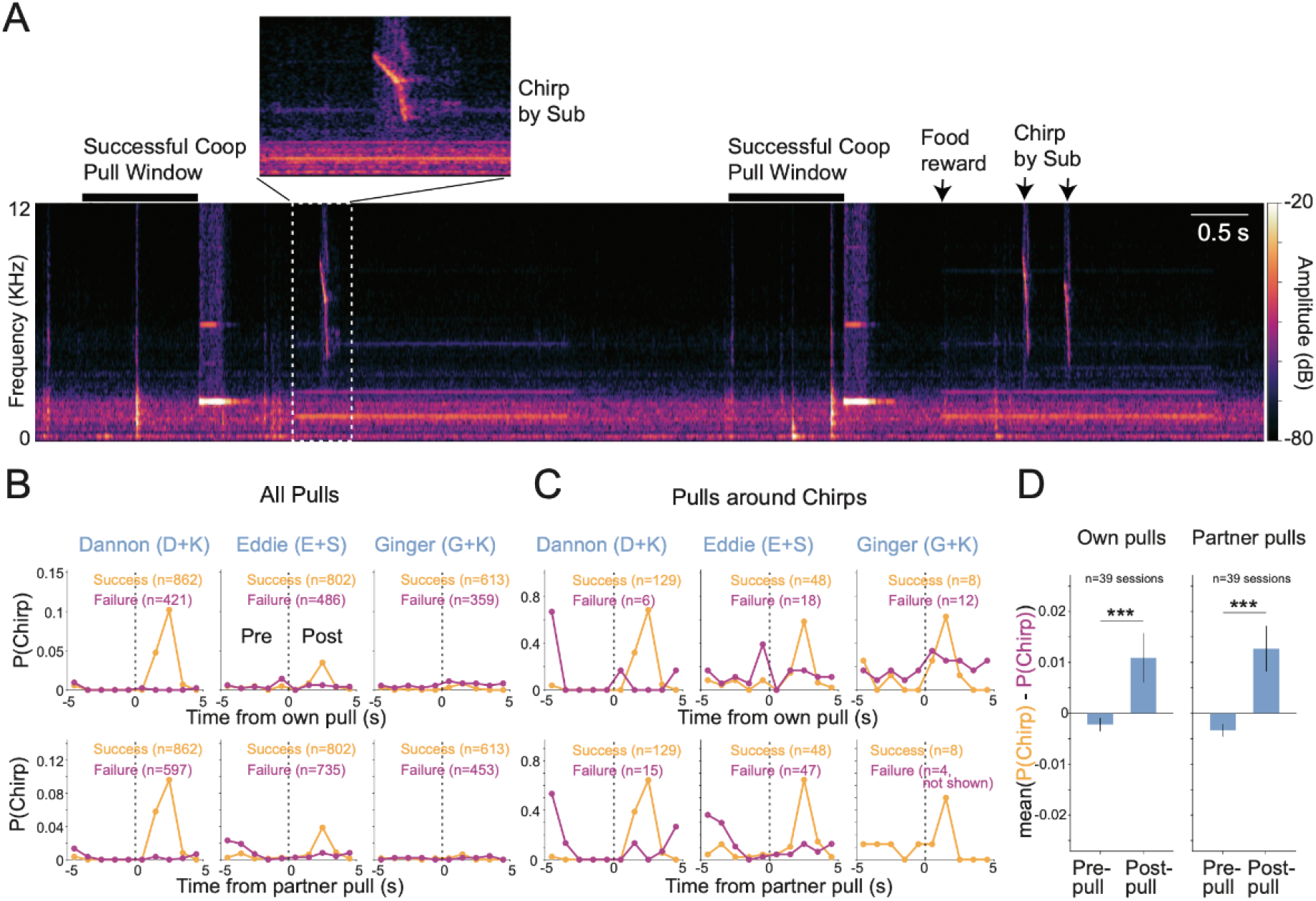
Subordinate marmoset’s chirps around cooperative pulls. (A) Example spectrogram of a subordinate monkey’s audio. It shows two successful cooperative pull windows (black bars), each followed by chirps from the subordinate (dashed box, enlarged in the inset, plus arrows). The color scale shows amplitude in dB. (B) Probability that the subordinate monkey gives at least one chirp in each 1 s bin, from 5 s before to 5 s after a pull. Successful pulls are orange and failed pulls are magenta. The top row is aligned to the subordinate’s own pulls and the bottom row to the partner’s pulls. All pulls from labeled sessions are included; n is the number of pulls. (C) Same as (B), but counting only pulls with at least one of the subordinate’s chirps within ±5 s. Curves based on 5 or fewer pulls are not drawn (Ginger, failed partner pulls, n = 4). (D) Mean of P(chirp | success) − P(chirp | failure) over the five pre-pull and five post-pull bins, for own pulls (left) and partner pulls (right). Bars show mean ± SEM across sessions (n = 39 sessions, all three dyads pooled). Pre vs. post was compared with a paired Wilcoxon signed-rank test (own pulls p = 3.1 × 104; partner pulls p = 1.3 × 104). ^***^ p < 0.001.

Across all pulls, the chirp probability peaked 1–2 s after successful cooperative pulls but remained near zero after failed cooperative attempts (Figure 3B). This pattern was observed regardless of whether the pulls were aligned to the subordinate’s own pulls or to the partners’. The peak probability was variable across animals and its ordering matches the dyads’ overall chirp rates (Figure 1E). Among the pulls that accompanied at least one subordinate chirp within ±5 s of the pulls, this pattern of chirps following the successful pulls became very clear in all three subordinates (Figure 3C). In contrast, chirps around the failed pulls occurred much less frequently and were not particularly associated with the time of pulls.

When we compared the average difference in chirp probability conditioned on success or failure outcome, it was near zero prior to the pull and was positive after the pull (Figure 3D), indicating a greater proportion of chirps following successful pulls. This pattern was observed around the subordinate’s own pulls (pre ∼−0.002, post ∼+0.011; p = 3.1 × 10^−4^, paired Wilcoxon signed-rank test) and around the partner’s pulls (pre ∼−0.003, post ∼+0.013; p = 1.3 × 10^−4^). Subordinate chirps therefore marked successful coordination, the mirror image of the dominants’ t-peeps that signified failed cooperation.

### Vocalization correlated with leadership in cooperation irrespective of dominance

Our prior work has found that the process of learning to cooperate shapes the relationship between how dominant animals and subordinate animals pull to achieve successful cooperation (Meisner et al). Here we asked how the leader-follower relationship of each partner changed due to training. For each successful cooperation event, we assigned the animal whose pull came first as the leader. For two out of three dyads, the leader role of the dominant marmosets decreased from the training to the trained stage, while this role increased for the subordinate marmosets (Figure 4A). We did not observe a significant change in leadership across the two stages for the third dyad. Thus, at least in two dyads, mastering the task resulted in a change in which partner led the lever-pulling, but not in a direction as expected with respect to social hierarchy. Leveraging on these changes in leader-follower relationship from training, we next examined if t-peeps and chirps, which were differently associated with social rank and cooperation outcome, had correlates of these changes.

**Figure 4.**
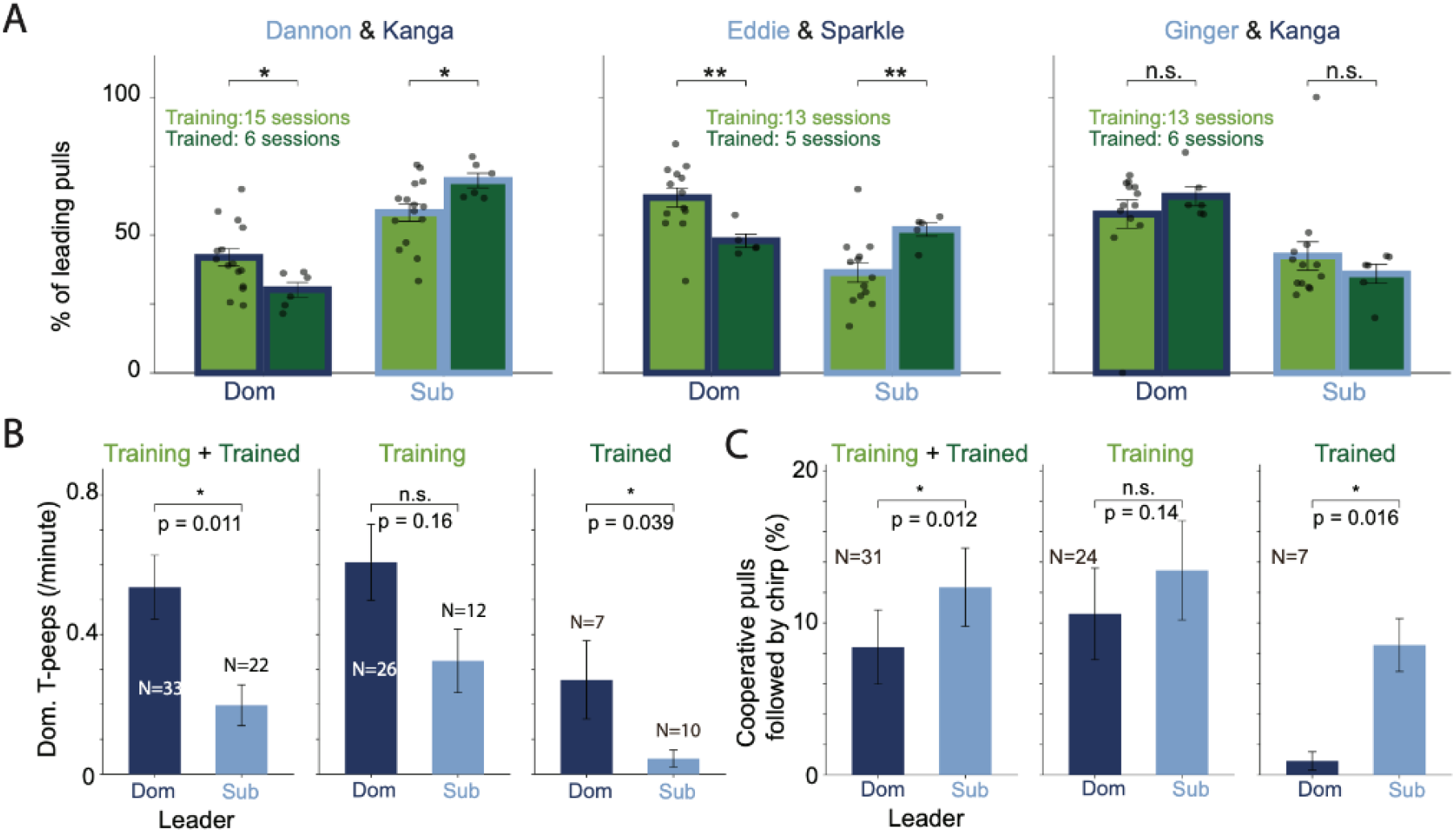
Leadership role in cooperation and its relation to vocalization. (A) Percentage of cooperative pulls led by the dominant (Dom) and subordinate (Sub) monkey in each dyad, during Training and Trained periods. The leader is the monkey whose successful pull came first. Dots show individual sessions. Mann–Whitney U test, Training vs. Trained. (B) T-peep vocalization per minute by session leader (either socially dominant or subordinate) with dyads pooled (Mann-Whitney U test).Columns show Training + Trained, Training only and Trained only. (C) Percentage of cooperative pulls followed by a chirp within 5 s, by leader (Wilcoxon signed-rank test). Columns show Training + Trained, Training only and Trained only. N = number of sessions. Bars show mean ± SEM. ^*^p < 0.05, ^**^p < 0.01; n.s., not significant.

#### Dominant t-peeps accompany dominant leadership in a session

Because dominant t-peeps were not closely tied to successful cooperative events (Figure 2B,C), we asked whether dominants t-peeped more in sessions they led. A session counted as ‘dominant-led’ if the dominant led more than half of its cooperative pulls, and as ‘subordinate-led’ otherwise. Across all sessions, dominant marmosets produced t-peeps more than twice as often in sessions they led (∼0.54 vs. ∼0.20 t-peeps/min; n = 33 vs. 22 sessions; p = 0.011, Mann–Whitney U test; Figure 4B). These t-peep patterns were most clear in the trained stage (∼0.27 vs. ∼0.04 t-peeps/min; n = 7 vs. 10; p = 0.039). The same trend was observed but was not significant during the training stage (∼0.61 vs. ∼0.32 t-peeps/min; n = 26 vs. 12; p = 0.16).

#### Subordinate chirps follow cooperative pulls they lead

We then asked whether a chirp from subordinate marmoset following a successful cooperative pull depended on who had led the pull. As mentioned above, all chirps within 5 s of a successful cooperative pull came from the subordinate marmoset(192 of 192). Across all sessions, chirps followed ∼12% of subordinate-led pulls and ∼8% of dominant-led pulls. The rate was higher for subordinate-led pulls in 23 of 31 sessions (p = 0.012, paired Wilcoxon signed-rank test; Figure 4C). During the training stage, the difference was smaller and not significant (∼13% vs. ∼11%; higher for subordinate-led pulls in 16 of 24 sessions; p = 0.14). However, once the dyads were trained, this effect became clear, where subordinates chirped almost only after cooperations they had led (∼9% vs. ∼1%; higher for subordinate-led pulls in all 7 sessions; p = 0.016). Together, these results link each partner’s call to its own role in the cooperation: dominant t-peeps were associated with dominant leadership, and subordinate chirps with subordinate leadership. Both of these associations became more reliable once the task was fully learned in the Trained stage.

### T-peeps and Chirps track opposite phases of cooperation

Given the relatively weak association of t-peeps from dominant animals around successful pulls compared to failed ones (Figure 2B), we hypothesized that they signalled a period of failed cooperation attempts. Thus, we next asked whether the use of these calls also depended on the state of cooperation. To answer this question, we divided each session into a series of cooperative bouts and the lulls between them, based on the naturally occurring gaps between successive cooperation rewards (Fig. 5A). Rewards came frequently, in general: 95% of gaps were 30 s or shorter, and the rest marked pauses in cooperative engagement.

**Figure 5.**
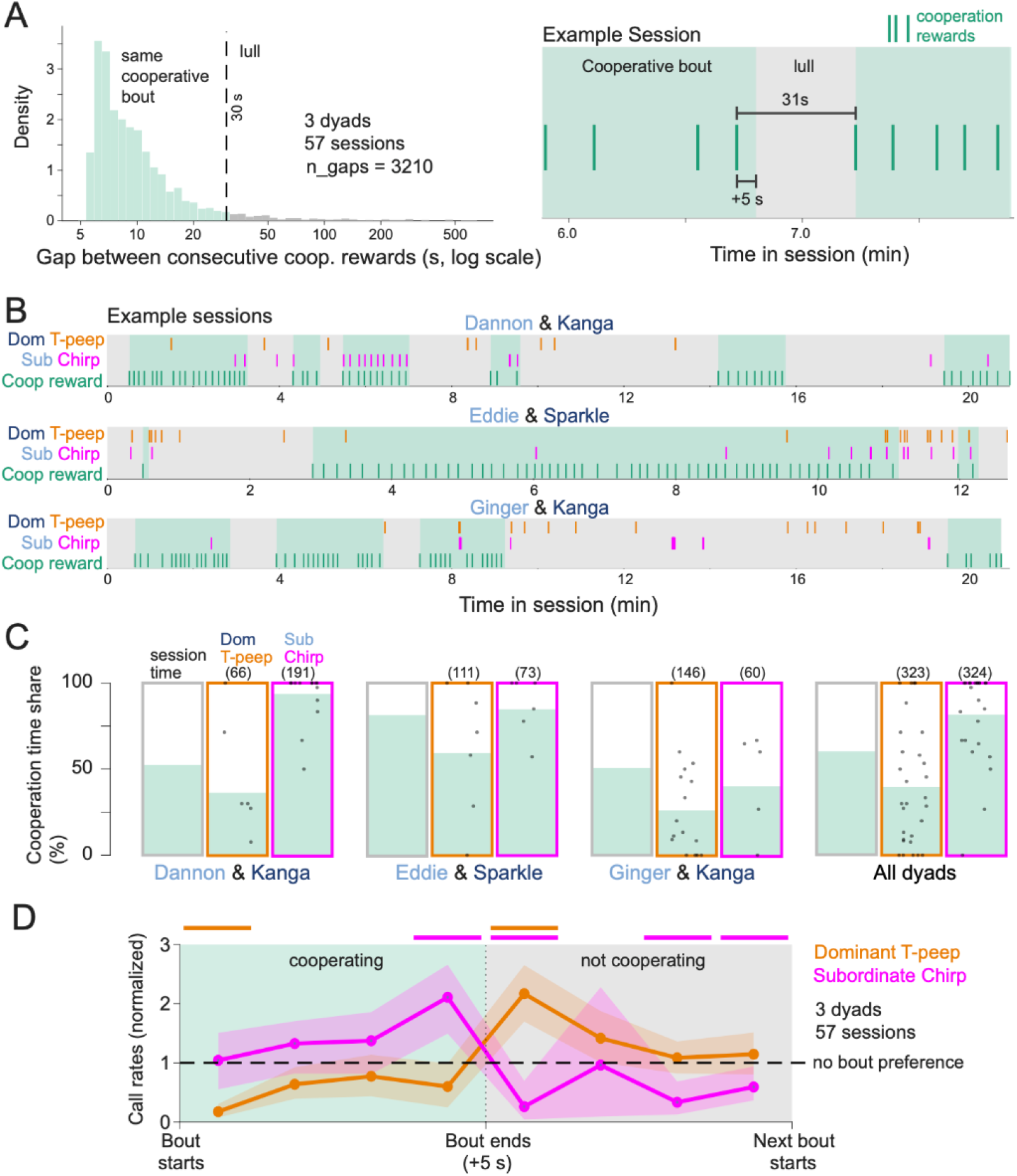
Calls relative to cooperative bouts and lulls. (A) Identification of cooperative bouts. Left: distribution of gaps between consecutive cooperation rewards, pooled across sessions and dyads. Gaps of 30 s or less (5% of gaps) count as part of the same bout, and longer gaps count as lulls. The 30 s cutoff minimized fragmented cooperation bouts. Right: an excerpt from one session illustrating this rule. Each bout runs from its first reward to 5 s after its last reward, so a chirp in response to the final reward still falls inside the bout. (B) One example session per dyad, with cooperative bouts shaded. (C) Share of task time spent cooperating (green portion), compared with the share of each call type made during cooperation (green portion). Calls are pooled across sessions, and numbers in parentheses are call counts. Dots show sessions with at least 3 calls of that type. (D) Call rates across one bout-to-lull cycle. Each bout and the lull after it are stretched to the same width (4 bins per part). Each rate is divided by the average of that call type’s bout rate and lull rate, with the two states weighted equally. Shading shows the 95% confidence interval from resampling sessions. Bars above the plot mark bins differing from 1 after Benjamini-Hochberg False Discovery Rate (q = 0.05, 8 bins per call type).

We found that subordinate chirps clustered inside the cooperative bouts, while dominant t-peeps often occurred in the lulls and just before bouts resumed (Fig. 5B). Across all dyads, the animals cooperated for 60.4% of task time (Figure 5C). Subordinate chirps were strongly over-represented during the cooperation bouts, with 81.8% of chirps falling inside the bouts. This chirp enrichment was present for two out of the three dyads (D-K: 93.7% of cooperation that happened 52.5% of time, E-S: 84.9% of cooperation that happened 81.4% of time, G-K: 40.0% of cooperation that happened 50.7% time). By contrast, dominant t-peeps were under-represented, with only 39.6% falling inside the bouts (Fig. 5C). In every dyad, t-peeps fell inside the cooperation bouts less often than expected from time spent cooperating (36.4% versus 52.5% for D-K, 59.5% versus 81.4% for E-S, and 26.0% versus 50.7% for G-K).

To further understand the relationship between these calls and the state of cooperation, we normalized the time of each cooperation bout and the following lull, and found that the two call types followed opposite time courses (Fig. 5D). Dominant t-peeps were nearly absent at the start of a cooperation bout (normalized rate 0.17, 95% CI 0.07–0.30) and stayed below the average session rate for the rest of the bout. Notably, they peaked right after the bout ended, in the first quarter of the lull (2.17, 95% CI 1.67–2.65), and then declined toward the average session rate as the lull continued. Both the decrease at the bout-onset and the increase at the lull-onset were significant (Fig. 5D). Again, the pattern for subordinate chirps was exactly the opposite of dominant t-peeps. Their rate increased across the cooperative bout and peaked in its final quarter (normalized rate 2.11, 95% CI 1.50–2.67). The chirp patterns dropped sharply once cooperation stopped (normalized rate 0.26, 95% CI 0.03–0.65) in the first quarter of the lull, and remained below the average session rate for most of the lull. Together, these results show that subordinate chirps track ongoing successful cooperation, while dominant t-peeps occur mainly when cooperation breaks down. This pattern suggests that dominant t-peeps act to re-engage the partner after a lull. Overall, we found multiple pieces of evidence that support divergent functions between dominant t-peeps and subordinate chirps for different functional aspects of cooperative interactions.

## DISCUSSION

Our results reveal a striking division of vocal communication in cooperating marmosets. Rather than using a common vocal signal to coordinate their actions, marmoset partners adopted complementary, rank-dependent signaling strategies. These strategies were selectively coupled to the outcome of their joint behavior required for cooperation success, to their own role in initiating it, and to whether cooperation was ongoing or had broken down. Dominant animals produced t-peeps after failed cooperative attempts, most often just as a bout of successful cooperation ended. By contrast, subordinate animals produced chirps after successful coordination, at rates that built up over sustained cooperation. Failed and successful cooperation were therefore signaled by different members of the dyad by way of their respective vocal calls. This vocalization schema was not simply a consequence of pulling or of general vocal activity. Both types of vocal signals were specifically associated with cooperative outcomes, dominant t-peeps declined markedly once cooperation was learned, and each call was linked to the cooperative event that the caller itself had led. Together, these findings suggest that vocal communication during cooperation is structured jointly by social hierarchy and task role, with partners occupying distinct communicative roles that may support both the acquisition and the maintenance of coordinated behavior.

### A functional role for vocalization around cooperative pulls

#### T-peeps

Our finding that dominant t-peeps follow failed cooperative pulls parallels the dominant marmosets’ gaze behavior we previously reported^10^: dominant animals made more social gaze than subordinates, and the amount of social gaze was correlated with the number of failed, not successful, pulls. Dominant t-peeps after failures may thus serve the same role as that of gaze: re-establishing coordination with the partner after a miss.

#### Chirps

In this study, we identified chirps purely by their acoustic marker. They occurred primarily after successful cooperative pulls, which were followed by liquid food rewards. This is in agreement with prior functional descriptions of marmoset calls with this acoustic signature as food calls^13^.

#### Other Calls

The low rates of phee and trills that we observed during the mutual cooperation task is not surprising. Trills are used between nearby animals but are not tied to any specific event, while phees are mainly contact calls produced when partners are out of view^16^. In our task, the partners could always see each other. Our prior work likewise found that phees increased only when an opaque barrier blocked vision, and occurred mostly during rest rather than active pulling^10^.

### Vocal signals follow the caller’s role in cooperation

Beyond social rank, each call type was linked to the caller’s own leadership of the cooperation. Dominants produced more t-peeps in sessions they led, and subordinates chirped mainly after cooperative pulls they had initiated, and almost exclusively so once the task was learned.

Although the increase in t-peeps by dominant animals when leading the pull appears to conflict with the use of t-pees with failure, the two are likely related. The leader of a cooperation is operationally defined as the animal that pulls first. An animal that tends to pull first is also the one more likely to pull without being reciprocated, and therefore experiencing failed pulls may enhance the need to recruit its partner. T-peeps may thus be produced by the partner that drives the cooperation and must wait for the other to join. Similarly, a subordinate that initiates a successful cooperation has, in a sense, achieved the outcome through its own action, and it is after these cooperations that it chirps.

In summary, our results suggest that although social rank determines which call an animal uses, its moment-to-moment role in the task determines when that call is produced. This dissociation between social role and task role echoes the observation that leadership changed with learning but not in a consistent direction relative to the hierarchy. Social rank and task role are separable dimensions of the partnership, and our findings support that vocal behavior reflects both. Notably, both vocal–leadership associations became significant after learning, which suggests that communicative roles may become more tightly bound to task roles as dyads converge on a stable coordination strategy.

### Communication across learning

The t-peep and chirp calls followed different trajectories across learning. Dominant t-peeps declined sharply once dyads were trained, whereas subordinate chirps fell only modestly and not significantly. This divergence suggests that the two vocal signals serve different stages of the cooperative interaction. A failure-related signal is most useful while partners are still discovering how to coordinate and becomes unnecessary once coordination is reliable. On the other hand, an outcome-related signal remains relevant whenever cooperation succeeds. This distinction has also been observed in human joint action, where explicit communication is most prominent during the acquisition of a joint task and gives way to implicit coordination as partners become practiced^19,20^. Training also changed leadership, shifting the dominant’s share of cooperation initiation in two of the three dyads. The decrease in dominant t-peeps may therefore partly reflect the dominant animal taking a less initiating role after learning.

### Parallels with human coordinative communication

The complementary organization we observed suggests parallels with how humans maintain coordination in joint action and conversation. When communication or coordination fails, human interlocutors initiate repair with short, specialized signals that recruit the partner to fix the problem^21,22^. When coordination succeeds, they produce brief acknowledgments that establish common ground^23^. The parallel with marmoset calls indicates that dividing communication labor between failure-related and success-related signals may be a general solution to the problem of coordinating joint action, and one that is already present in a cooperatively breeding primate. This fits with proposals that cooperative breeding promoted the social-interactional foundations of communication^5,6^. Vocal coordination during cooperative tasks has also been reported in other species, such as bottlenose dolphins^24^, suggesting role-dependent signaling during joint action as a strategy that extends beyond primates.

In contrast to a stable rate of social gaze across training^10^, a significant reduction in the rate of t-peeps after training suggests that they are mainly a strategy for *learning* and not *learned* cooperation. The significant increase in the rate of t-peeps at the onset of cooperative lulls suggests that they are not pull-by-pull but a slower timescale coordination strategy. The overall fewer occurrences of t-peeps compared to social gaze events during cooperation could reflect a fundamental difference in the salience of signalling modalities. An auditory signal is detectable by a partner that may or may not be engaged in visual monitoring, and infrequent usage may further enhance its salience. T-peep usage in our data thus suggests that vocal signaling during cooperation does not require the frequency of non-verbal signaling such as gaze or gesture. This parallels observation in humans, where non-verbal communication often carries much of the coordination load in joint action, and that speech can be unnecessary or even costly^25–27^.

### .Dyad and relationship differences

Although the rank-dependent division of calls was observed in all three dyads, its magnitude varied with the relationship between partners. The female–female, parent–offspring dyad (Ginger–Kanga) produced the most dominant t-peeps and the fewest subordinate chirps. It was also the only dyad in which chirps were not concentrated within cooperative bouts. On the other hand, the male–female pair-bonded dyads showed the strongest coupling of chirps to successful cooperation. These differences may reflect the nature of the relationship: kinship versus pair-bond, female–female versus male–female, and the size of the age and rank gap between partners. In our prior work^10,28^, we showed that kinship and sex shape cooperative strategies, and our current results suggest these social factors may also guide the vocal behavior during cooperation. Testing this will require more dyads that systematically vary relationship type.

### Limitations and future directions

Several limitations exist in our work. First, the sample size was small: three dyads, five individuals and six dyadic roles. One marmoset (Kanga) was the dominant partner in two of the three dyads, and in both male–female pairs the dominant animal was the female. Social rank is therefore partly confounded with sex and with individual identity. Each dyad does show the same rank-dependent division of calls with a different subordinate partner, and this pattern was observed in a female–female dyad, which argues against a pure sex effect. Still, dyads in which a male is dominant, or in which the same individual holds different ranks with different partners, would be needed to systematically separate rank from sex and identity. Second, sessions were pooled across dyads for several analyses. Future analyses with mixed-effects models that include dyad and individual as random effects will be important to confirm that the effects are not driven by between-dyad differences. Finally, all of our findings are correlational. Whether t-peeps and chirps alter the partner’s behavior, and thus serve as true communicative signals rather than by-products of the caller’s state, remains to be tested. Analyses of partner pulling and gaze conditioned on preceding calls, playback experiments, and integration of vocal and gaze data in the same sessions will help resolve this.

Together, our findings suggest that vocal signaling during cooperative interaction is not necessarily shared symmetrically between partners. Instead, it can be organized by hierarchical role and by task role: different individuals signal complementary information about failure and success, and the timing of those signals follows each partner’s contribution to the joint action and the waxing and waning of cooperation itself. Such a division of communicative labor may help partners discover and maintain coordination, and it identifies the marmoset as a powerful model for studying how social structure shapes communication during joint behavior.

## ACKNOWLEDGMENTS

This work was supported by the National Institute of Mental Health (R21 MH126072 and RF1 MH138396, SWCC, ASN, MPJ; K99 MH142687, WS), the National Institute on Drug Abuse (U01 DA063534, SWCC, ASN, MPJ), the National Science Foundation Graduate Research Fellowship (DGE2139841, OCM), the Simons Foundation Autism Research Initiative (SFARI 875855, SWCC, ASN, MPJ), Wu Tsai Institute at Yale University (SWCC, ASN, MPJ, WS), and a National Eye Institute core grant for vision research (P30 EY026878, Yale University). We thank the veterinary and husbandry staff at Yale for their excellent animal care.

## AUTHOR CONTRIBUTIONS

Conceptualization: GN, SWCC, ASN, MPJ; Experimental Design: OCM, WS, SWCC, ASN; Data Collection: GN, OCM, WS; Data Analysis: GN; Supervision: SWCC, ASN, MPJ; Writing: GN, SWCC, ASN, MPJ.

## METHODS

### Subjects

Three dyads of familiar adult common marmosets (*Callithrix jacchus*) were tested: two male–female pair-bonded dyads (Eddie–Sparkle and Dannon–Kanga) and one female–female parent–offspring dyad (Ginger–Kanga). Kanga took part in two dyads, giving five individual animals and six dyad roles.

### Automated Apparatus

Cooperative pulling was assessed with MarmoAAP, a fully automated apparatus^11^. It consists of two adjacent transparent boxes, each with a lever and a liquid reward tube at the front slot. An overhead GoPro camera recorded video, and a microphone (Qzt Electronic) in the back corner of each box recorded audio.

### Behavioral Task

Marmosets were trained to pull a lever, first independently and then cooperatively, for a liquid reward (6 g of marshmallow fluff per 20 mL of water). Sessions lasted 20–30 min, five days per week. Animals had no access to food or water for 1–3 h before each session. Training began with a self-reward stage, in which each animal pulled to earn its own reward. Cooperation training followed, with the window shrinking from 3 s to 2 s, 1.5 s and finally 1 s: both animals had to pull within that window of each other to be rewarded. Apparatus and training procedures otherwise followed Meisner et al. (2024). All analyses used the Mutual Cooperation (MC) task at the 1 s threshold, both for the *Training* and *Trained* stages. A pull was registered when the lever passed a set position threshold, and the task-control computer recorded its time at 10 kHz. The lever had to be released after crossing the threshold for the pull to count. When both animals pulled within 1 s of each other, a 2 kHz tone sounded and both received 0.2 mL of reward. A pull not matched by the partner within 1 s was a failed pull. The task had no imposed trial structure; animals made a continuous series of self-paced pulls. Successful pulls were the two pulls (one per animal) just before a reward delivered to both levers at once. All other pulls were classed as failed. Lever-pulling and vocal behaviors of each dyad were examined at two stages.The Training phase comprised the sessions conducted during MC 1 s training, totaling 42 sessions across the three dyads (Dannon–Kanga, n=15 sessions; Eddie–Sparkle, 13; Ginger–Kanga, 14). The Trained phase comprised the later sessions conducted after each dyad had reached either 50% cooperative success or had performed at least 4 sessions of the condition^10^, totaling 17 sessions (Dannon–Kanga, 6; Eddie–Sparkle, 5; Ginger–Kanga, 6). Unless stated otherwise, analyses used Training sessions.

### Dominance Hierarchy Assignment

In male–female pairs the female was assigned as dominant; in the female–female dyad the older female was assigned as dominant. Feeding order at daily meals and preferred resting spots confirmed these assignments^10^. The dominant animals were Sparkle (over Eddie), Kanga (over Dannon) and Kanga (over Ginger).

### Vocalization Analysis

Audio was labeled manually using *Audacity* software. The two microphone tracks of a session were aligned to session start, and each call was assigned to the animal whose microphone recorded it at higher amplitude. Each call’s type was identified from established marmoset call categories (REF), and its start and end times were marked. Four call types were labeled: chirp, t-peep, trill and phee. For sessions without microphone recordings, audio was extracted from the video cameras and labeled the same way.

#### Calls per session

For each animal and session, we counted the calls of each type. Dominant and subordinate counts were compared within the same Training sessions, with dyads pooled (Fig. 1C). Training and Trained counts were compared within each dyad, for the dominant animal’s t-peeps (Fig. 1D) and the subordinate animal’s chirps (Fig. 1E).

#### Call probability around pulls

We split the window from −5 s to +5 s around each pull into ten 1 s bins. Each bin was scored by whether it held at least one call onset by the calling animal. The call probability in a bin was the fraction of pulls with a call in that bin, computed separately for successful and failed pulls. Only pulls from sessions in which the caller’s audio was labeled were included.

#### Pre-vs. post-pull effect of success

For each session, we took P(call | success) − P(call | failure) in each bin. We then averaged it over the five pre-pull bins and over the five post-pull bins, for own and partner pulls (Figs. 2D, 3D). Only sessions with more than 5 successful and more than 5 failed pulls were included (n = 39 sessions).

#### Statistical Analysis

Summary statistics are reported as mean ± SEM across sessions, and all tests were two-sided with significance at p < 0.05.

### Analysis of Leadership of cooperative pulls

For each successful cooperative pull, each animal’s successful pull was its last pull since the previous reward. The animal whose successful pull came first was the leader of that cooperation, and rewards in which both pulls had the same timestamp were excluded. For each session, we computed the percentage of successful cooperative pulls led by each animal, so that the two values sum to 100% (Fig. 4A).

#### Dominant t-peeps by session leader

Sessions were classed as dominant-led if the dominant led more than 50% of the session’s successful cooperative pulls, and as subordinate-led if it led less than 50%. Sessions at exactly 50%, or with fewer than 10 successful cooperative pulls, were excluded, leaving 55 sessions. The dominant t-peep rate was the number of dominant t-peep onsets between the session’s first and last lever event, divided by that span in minutes. Only the dominant animal’s t-peeps were used, and dyads were pooled (Fig. 4B).

#### Chirps after cooperative pulls by leader

For each successful cooperative pull, we scored whether a chirp onset occurred within 5 s after the later of the two successful pulls, which completes the cooperation. For each session, we then computed the percentage of dominant-led pulls and of subordinate-led pulls that were followed by a chirp. Only sessions with at least one chirp after a successful cooperative pull were included (n = 31 sessions across both stages, 24 Training and 7 Trained), and dyads were pooled (Fig. 4).

#### Statistical Analysis

For leadership, the per-session percentage of dominant-led pulls was compared between Training and Trained sessions within each dyad using Mann–Whitney U tests (Fig. 4A). Dominant t-peep rates were compared between dominant-led and subordinate-led sessions using Mann–Whitney U tests, separately for all sessions, Training sessions and Trained sessions (Fig. 4B). Per-session chirp percentages after subordinate-led and dominant-led pulls were compared using paired Wilcoxon signed-rank tests (Fig. 4C).

### Analysis of Cooperation bouts and lulls

We analyzed all mutual-cooperation sessions at the 1 s pull threshold that had labeled vocalizations. While the task did not have an inherent trial structure, the dyads worked to maintain consecutive cooperation rewards at least about 5 s apart^10,28^. The distribution of gaps between consecutive rewards (n = 3,210; median 8.4 s) had a single mode near 7 s and a long right tail, with no clear valley separating gaps within a bout from pauses between bouts (Fig. 5A, left). To choose a cutoff, we fit a three-component Gaussian mixture model to the log10 gap durations, pooling all sessions. The components were centered at 6.7 s (41% of gaps), 10.6 s (48%) and 31.9 s (11%). The posterior probability of the slowest component passed 0.5 at 20.6 s. We used a more conservative cutoff of 30 s, where that posterior is about 0.99. Gaps longer than 30 s were classified as lulls (5.2% of gaps). Runs of rewards separated by gaps of 30 s or less formed cooperative bouts, and a single isolated reward counted as a one-reward bout. Each bout ran from its first reward to 5 s after its last reward (Fig. 5A, right). This 5 s extension matches the window in which subordinate chirps followed a successful pull, so a chirp given in response to a bout’s final reward still fell inside the bout. All remaining task time was classified as not cooperating. This included lulls between bouts, the period before the first reward and the period after the last bout.

#### Share of calls during cooperation

For each dyad and for all dyads pooled, we computed three shares, pooling time and calls across sessions: the share of task time spent cooperating, the share of dominant t-peeps produced while cooperating, and the share of subordinate chirps produced while cooperating (Fig. 5C). A call type was considered enriched during cooperation if its share was larger than the time share, and depleted if its share was smaller. Session-level shares are shown for sessions with at least three calls of the given type.

#### Call rates across the bout–lull cycle

Each bout was paired with the stretch that followed it. That stretch ran from the end of the bout (last reward plus 5 s) to the start of the next bout, or to the end of the task for a session’s last bout. We excluded the period before each session’s first bout, as well as any pair in which either part had zero duration. This left out 207 bout–lull pairs from 57 sessions. Each bout and each lull was divided into four bins of equal duration such that bouts and lulls of different lengths were placed on a common normalized time axis. Calls were counted and bin durations summed across all pairs and sessions, and a call rate was computed for each bin. Dyads spent different fractions of the cycle cooperating. To make rates comparable across dyads, each bin’s rate was divided by the unweighted mean of the pooled within-bout rate and the pooled within-lull rate such that the two states were weighted equally rather than by their durations. On this scale, a value of 1 means the same rate in bouts and lulls, and values of 2 or 0 mean that all calls of that type fell in one state. A time-weighted normalization capped the highest attainable within-bout value at the inverse of the cooperating fraction (for example, about 1.2 for Eddie–Sparkle, who cooperated for 81% of task time).

#### Statistical Analysis

Ninety-five percent confidence intervals came from resampling sessions with replacement 5,000 times and recomputing the normalized rates for each resample. For each bin, a two-sided bootstrap p-value was computed as twice the smaller of two fractions: resampled values at or below 1, and resampled values at or above 1. P-values were corrected with the Benjamini–Hochberg false discovery rate procedure (q = 0.05), applied separately to the eight bins of each call type.

### Software Tools

All statistical analyses were performed in Python (NumPy, pandas, SciPy, scikit-learn), and figures were drawn in MATLAB R2025a.

## Supplementary Information

**Chirps**: Each chirp is a downward sweep from about 10–12 kHz to 5–6 kHz, about 50 ms long.

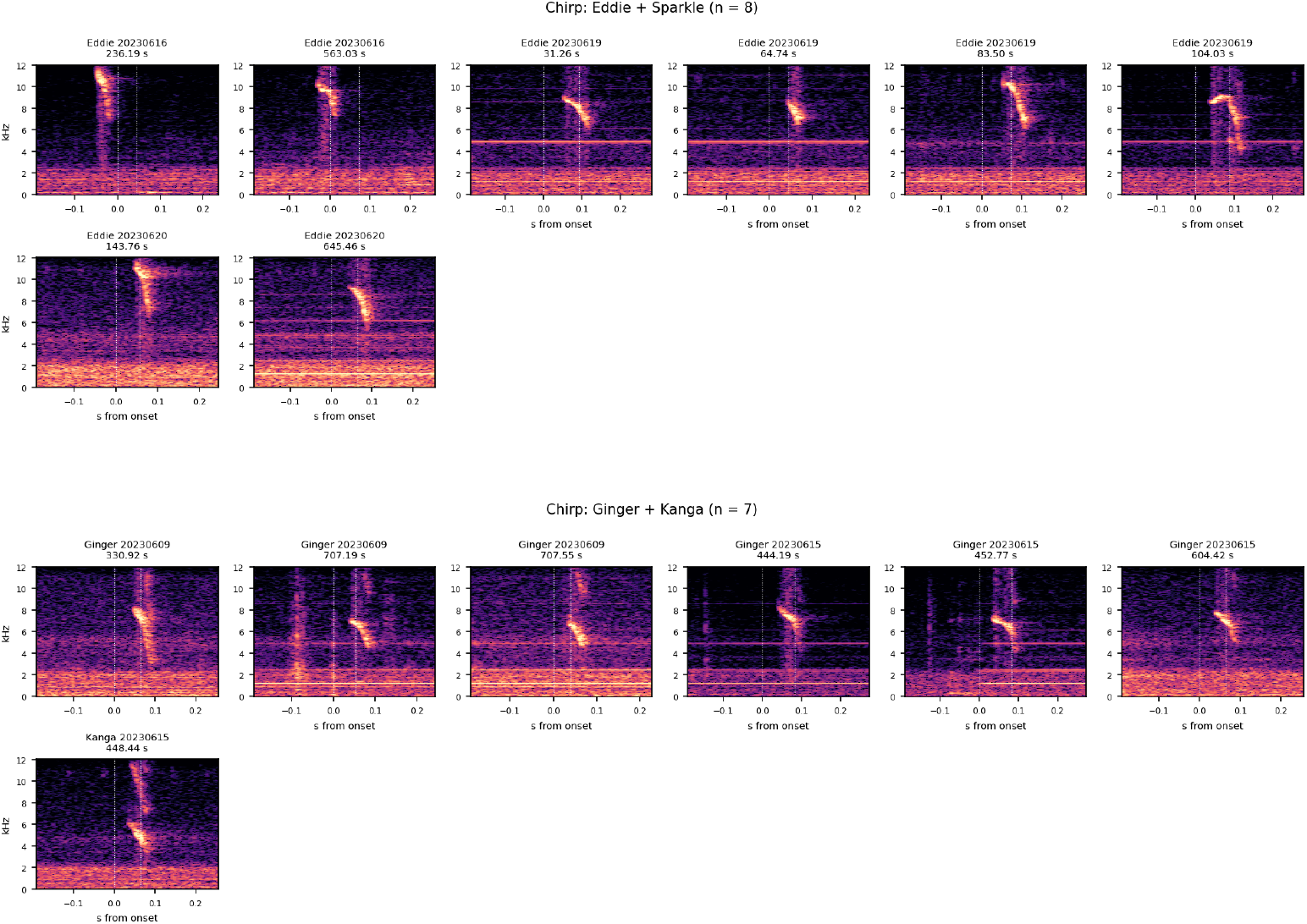

**T-peeps**: t-peeps are visible as a flat-to-wavering tone near 6 kHz, 60–120 ms long.

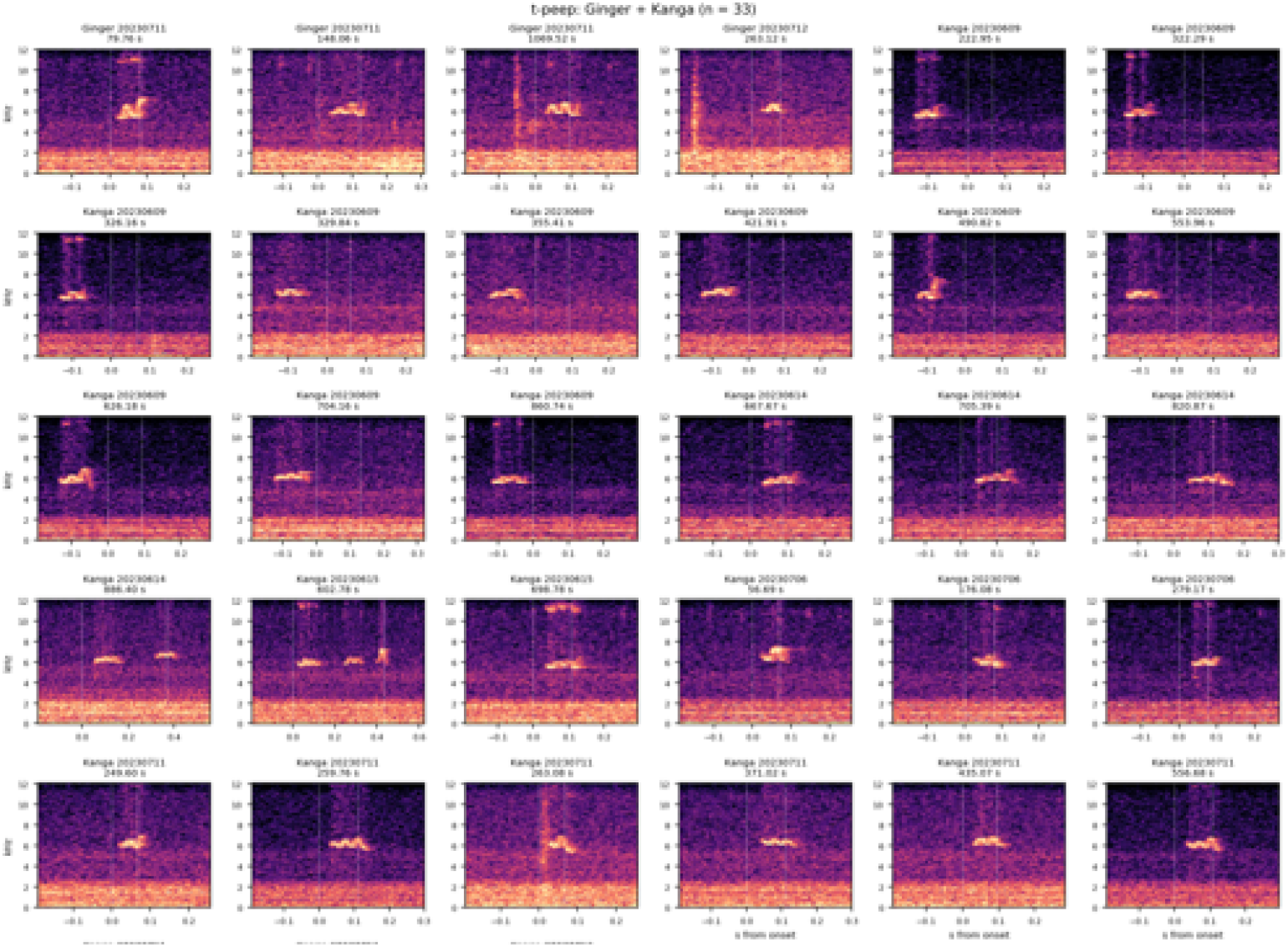

## References

1. Miller, C.T., Freiwald, W.A., Leopold, D.A., Mitchell, J.F., Silva, A.C., and Wang, X. (2016). Marmosets: a neuroscientific model of human social behavior. Neuron 90, 219–233. 10.1016/j.neuron.2016.03.018.

2. Burkart, J.M., Fehr, E., Efferson, C., and van Schaik, C.P. (2007). Other-regarding preferences in a non-human primate: common marmosets provision food altruistically. Proc. Natl. Acad. Sci. 104, 19762–19766. 10.1073/pnas.0710310104.

3. Takahashi, D.Y., Narayanan, D.Z., and Ghazanfar, A.A. (2013). Coupled oscillator dynamics of vocal turn-taking in monkeys. Curr. Biol. 23, 2162–2168. 10.1016/j.cub.2013.09.005.

4. Choi, J.Y., Takahashi, D.Y., and Ghazanfar, A.A. (2015). Cooperative vocal control in marmoset monkeys via vocal feedback. J. Neurophysiol. 114, 274–283. 10.1152/jn.00228.2015.

5. Burkart, J.M., Adriaense, J.E.C., Brügger, R.K., Miss, F.M., Wierucka, K., and van Schaik, C.P. (2022). A convergent interaction engine: vocal communication among marmoset monkeys. Philos. Trans. R. Soc. B 377, 20210098. 10.1098/rstb.2021.0098.

6. Takahashi, D.Y., Liao, D.A., and Ghazanfar, A.A. (2017). Vocal Learning via Social Reinforcement by Infant Marmoset Monkeys. Curr. Biol. 27, 1844–1852.e6. 10.1016/j.cub.2017.05.004.

7. Liao, D.A., Zhang, Y.S., Cai, L.X., and Ghazanfar, A.A. (2018). Internal states and extrinsic factors both determine monkey vocal production. Proc. Natl. Acad. Sci. 115, 3978–3983. 10.1073/pnas.1722426115.

8. Grijseels, D.M., Prendergast, B.J., Gorman, J.C., and Miller, C.T. (2023). The neurobiology of vocal communication in marmosets. Ann. N. Y. Acad. Sci. 1528, 13–28. 10.1111/nyas.15057.

9. Eliades, S.J., and Miller, C.T. (2017). Marmoset vocal communication: Behavior and neurobiology. Dev. Neurobiol. 77, 286–299. 10.1002/dneu.22464.

10. Meisner, O.C., Shi, W., Nair, A., Nandy, G., Jadi, M.P., Nandy, A.S., and Chang, S.W.C. (2025). Diverse and flexible strategies enable successful cooperation in marmoset dyads. Curr. Biol. 35, 4509–4521.e5. 10.1016/j.cub.2025.08.005.

11. Meisner, O.C., Shi, W., Fagan, N.A., Greenwood, J., Jadi, M.P., Nandy, A.S., and Chang, S.W. (2024). Development of a Marmoset Apparatus for Automated Pulling to study cooperative behaviors. eLife 13, RP97088. 10.7554/elife.97088.

12. Epple, G. (1968). Comparative Studies on Vocalization in Marmoset Monkeys (Hapalidae). Folia Primatol. (Basel) 8, 1–40. 10.1159/000155129.

13. Vitale, A., Zanzoni, M., Queyras, A., and Chiarotti, F. (2003). Degree of social contact affects the emission of food calls in the common marmoset (Callithrix jacchus). Am. J. Primatol. 59, 21–28. 10.1002/ajp.10060.

14. Turesson, H.K., Ribeiro, S., Pereira, D.R., Papa, J.P., and De Albuquerque, V.H.C. (2016). Machine Learning Algorithms for Automatic Classification of Marmoset Vocalizations. PLOS ONE 11, e0163041. 10.1371/journal.pone.0163041.

15. Bezerra, B.M., and Souto, A. (2008). Structure and usage of the vocal repertoire of Callithrix jacchus. Int. J. Primatol. 29, 671–701. 10.1007/s10764-008-9250-0.

16. Agamaite, J.A., Chang, C.-J., Osmanski, M.S., and Wang, X. (2015). A quantitative acoustic analysis of the vocal repertoire of the common marmoset (Callithrix jacchus). J. Acoust. Soc. Am. 138, 2906–2928. 10.1121/1.4934268.

17. Pomberger, T., Risueno-Segovia, C., Gultekin, Y.B., Dohmen, D., and Hage, S.R. (2019). Cognitive control of complex motor behavior in marmoset monkeys. Nat. Commun. 10, 3796. 10.1038/s41467-019-11714-8.

18. Digby, L.J. (1995). Social organization in a wild population of Callithrix jacchus: II. Intragroup social behavior. Primates 36, 361–375. 10.1007/BF02382859.

19. Sebanz, N., Bekkering, H., and Knoblich, G. (2006). Joint action: bodies and minds moving together. Trends Cogn. Sci. 10, 70–76. 10.1016/j.tics.2005.12.009.

20. Vesper, C., Butterfill, S., Knoblich, G., and Sebanz, N. (2010). A minimal architecture for joint action. Neural Netw. 23, 998–1003. 10.1016/j.neunet.2010.06.002.

21. Schegloff, E.A., Jefferson, G., and Sacks, H. (1977). The Preference for Self-Correction in the Organization of Repair in Conversation. Language 53, 361–382. 10.2307/413107.

22. Dingemanse, M., Roberts, S.G., Baranova, J., Blythe, J., Drew, P., Floyd, S., Gisladottir, R.S., Kendrick, K.H., Levinson, S.C., Manrique, E., et al. (2015). Universal Principles in the Repair of Communication Problems. PLOS ONE 10, e0136100. 10.1371/journal.pone.0136100.

23. Clark, H.H., and Brennan, S.E. (1991). Grounding in communication. In Perspectives on Socially Shared Cognition, L. B. Resnick, J. M. Levine, and S. D. Teasley, eds. (American Psychological Association), pp. 127–149. 10.1037/10096-006.

24. King, S.L., Guarino, E., Donegan, K., McMullen, C., and Jaakkola, K. (2021). Evidence that bottlenose dolphins can communicate with vocal signals to solve a cooperative task. R. Soc. Open Sci. 8, 202073. 10.1098/rsos.202073.

25. Pezzulo, G., Donnarumma, F., and Dindo, H. (2013). Human Sensorimotor Communication: A Theory of Signaling in Online Social Interactions. PLoS ONE 8, e79876. 10.1371/journal.pone.0079876.

26. Wyman, E., Rakoczy, H., and Tomasello, M. (2013). Non-verbal communication enables children’s coordination in a “Stag Hunt” game. Eur. J. Dev. Psychol. 10, 597–610. 10.1080/17405629.2012.726469.

27. Brennan, S.E., Chen, X., Dickinson, C.A., Neider, M.B., and Zelinsky, G.J. (2008). Coordinating cognition: The costs and benefits of shared gaze during collaborative search. Cognition 106, 1465–1477. 10.1016/j.cognition.2007.05.012.

28. Shi, W., Meisner, O.C., Jadi, M.P., Chang, S.W.C., and Nandy, A.S. (2026). Canonical decision computations underlie behavioral and neural signatures of cooperation in primates. Neuron 114, 2657–2667.e5. 10.1016/j.neuron.2026.04.022.

